# Investigating DNA methylation changes associated with food production using paleogenomes

**DOI:** 10.1101/2023.11.04.565610

**Authors:** Sevim Seda Çokoğlu, Dilek Koptekin, Fatma Rabia Fidan, Mehmet Somel

## Abstract

The Neolithic transition introduced major diet and lifestyle changes to human populations across continents. Beyond well-documented bioarchaeological and genetic effects, whether these changes also had molecular-level epigenetic repercussions in past human populations has been an open question. In fact, methylation signatures can be inferred from UDG-treated ancient DNA through postmortem damage patterns, but with low signal-to-noise ratios; it is thus unclear whether published paleogenomes would provide the necessary resolution to discover systematic effects of lifestyle and diet shifts. To address this we compiled UDG-treated shotgun genomes of 13 pre-Neolithic hunter-gatherer (HGs) and 21 Neolithic farmer (NFs) individuals from West and North Eurasia, published by six different laboratories and with coverage c.1x-58x (median=9x). We used epiPALEOMIX and a Monte Carlo normalization scheme to estimate methylation levels per genome. Our paleomethylome dataset showed expected genome-wide methylation patterns such as CpG island hypomethylation. However, analysing the data using various approaches did not yield any systematic signals for subsistence type, genetic sex, or tissue effects. Comparing the HG-NF methylation differences in our dataset with methylation differences between hunter-gatherers vs. farmers in modern-day Central Africa also did not yield consistent results. Meanwhile, paleomethylome profiles did cluster strongly by their laboratories of origin. Our results mark the importance of minimizing technical noise for capturing subtle biological signals from paleomethylomes.

## Introduction

The last 12,000 years saw diverse human populations shift from mobile hunter-gathering to Neolithic lifeways involving sedentism and food production. These Neolithic transitions not only brought about changes in diet but also major shifts in daily activities, an increase in population density, as well as institutionalized social inequalities (Bar-Yosef and Belfer-Cohen, 1992; Richards, 2002). Beyond their social impact, how these changes shaped human health, physiology, genetics and epigenetics has long been debated. Anthropological evidence points to negative outcomes related to dietary constraints and high population density, such as increasing prevalence of growth disruption, anaemia, or dental caries in archaeological human remains from Neolithic populations compared to foragers (Latham, 2013; Larsen, 2006). Meanwhile, population genomic studies have reported multiple loci that evolved under positive selection pressures related to agriculture and pastoralism. These include the FADS genes involved in polyunsaturated fatty acid metabolism (Buckley *et al*., 2017) and the LCT gene responsible for lactase persistence (Tishkoff *et al*., 2007). Even though these selection pressures appear to have gained strength multiple millennia later than the original transitions to food production (Burger *et al*., 2020; Mathieson and Mathieson, 2018), their documentation is consistent with the notion that food production had significant long-term impacts on human physiology.

It might be likewise expected that Neolithic transitions shifted human epigenetic profiles. Indeed, changes in overall methylation levels have been found in leukocytes related to vegetable-rich versus fat- and meat-rich diets in a human sample from the USA (Zhang *et al*., 2011). Even more relevant are the results by Fagny and colleagues, who compared blood methylation profiles between present-day hunter-gatherers (MHGs) and farmers (MFs) living in central Africa (Fagny *et al*., 2015). These authors reported thousands of loci showing differential methylation patterns correlated with both historical and recent shifts in lifestyle; they further associated these changes with immune- and development-related pathways. Whether similar past Neolithic human populations experienced similar lifestyle- and diet-related epigenetic shifts has remained an open question.

Unfortunately, most epigenetic information related to physiology is lost in ancient specimens as soft tissue and RNA are not preserved (see Smith *et al*., 2019 for an exception). However, it has been shown that cytosine methylation sites can survive in ancient DNA. A number of studies have used standard protocols for methylation profiling, such as bisulphite sequencing and immunoprecipitation, on ancient DNA (Smith *et al*., 2015; Seguin-Orlando et al., 2015) (Llamas et al., 2012). Meanwhile, methylation information can also be indirectly inferred from sequencing data from ancient DNA molecules treated with the UDG (uracil-DNA glycosylase) enzyme. This is based on the knowledge that after death, aDNA molecules undergo widespread cytosine deamination at their broken ends, resulting in C→U (uracil) transitions if the cytosine is unmethylated, and in C→T (thymine) transitions if the cytosine is methylated (Briggs *et al*., 2007). Treatment of aDNA with UDG eliminates uracil nucleotides from DNA, and when such UDG-treated aDNA is shotgun sequenced, the level of observed C→T transitions at CpG sites allows inferring the relative methylation level at those loci (Pedersen *et al*., 2014).

Over the last decade, a growing number of studies have reported succesful retrieval of methylation patterns in past organisms using this approach (Orlando *et al*., 2015). In 2014, Pedersen and colleagues studied 20x coverage UDG-treated genomic data produced from a 4000-year-old hair sample from Greenland (Pedersen *et al*., 2014). These authors reported significant correlations between genome-wide methylation levels inferred from this data with methylation measured in present-day human tissues, with the highest correlations found with hair. This study also found expected signals of hypomethylation in CpG islands in the paleomethylome data and further inferred the age of the ancient individual using a methylation clock. The same year, studying the 52x-coverage Neanderthal and 30x-coverage Denisovan genomes derived from bone material, Gokhman and colleagues (Gokhman *et al*., 2014) found overall low CpG methylation rates (<1.5%) as inferred from postmortem deamination; however, binning those methylation scores yielded high correlations with global methylation patterns measured in modern-day human bone samples. These authors further used this data to predict a number of loci, developmental genes, that might be differentially methylated between archaic hominins and modern humans. In 2016, Hanghøj and colleagues published the epiPALEOMIX MethylMap algorithm for estimating methylation scores in UDG-treated ancient DNA libraries with sufficient (e.g. >2x) coverage (Hanghøj *et al*., 2016). Applying their algorithm to published ancient human genomes, these authors showed tissue-based clustering among at least some of the paleomethylomes they analysed. Successful retrieval of paleomethylation signatures has also been reported for other species, including barley, maize, and horses (Wagner *et al*., 2020; Smith et al., 2014; Liu et al., 2023).

Therefore, despite the promising results described above, whether lifestyle-related paleomethylation signatures may be retrievable from ancient bone and tooth material is unknown. It is also unclear whether paleomethylome profiles inferred from data originating from different labs and different coverages could be easily comparable. This is a particularly challenging task because paleomethylome profiles are inferred indirectly, depending on the presence of random postmortem damage at read ends. The signal-to-noise ratio per locus is hence much lower compared to information collected using bisulphite sequencing on present-day tissue samples. Therefore the technical noise caused by different lab protocols could readily overshadow biological signals.

Here we address these issues by investigating systematic methylation differences in 34 published paleogenomes from hunter-gatherer (HG) and Neolithic farmer (NF) contexts, produced by different laboratories and with a range of depth-of-coverages. We further ask whether convergent HG-NF epigenetic shifts can be detected between ancient and present-day populations.

## Results

Our dataset comprises published paleogenomes of 13 HGs (45kya-4kya) and of 21 NFs (8.5kya-5kya) from Eurasia, all shotgun-sequenced and UDG-treated, and originating from 6 different laboratories and 8 different publications (Kılınç *et al*., 2021; Antonio et al., 2019; Fu et al., 2014; Günther et al., 2018; Lazaridis et al., 2014; Marchi et al., 2022; Sánchez-Quinto et al., 2019; Seguin-Orlando et al., 2014) (Supplementary Table 1; SupplementaryF igures 1-2). As Figure 1A shows, our sample was concentrated in West and North Eurasia to limit genetic background variation (Methods). Of the 34 genomes, 23 were derived from bone and 11 from tooth; 12 were female and 22 male; 4 were produced using single-stranded and the rest double-stranded library protocols. The genome coverages ranged from c.1x-58x (median=9x). Genomes from different publications had different coverage levels (ANOVA *P*=3E-14), but the subsistence type groups (HG vs NF) coverages were not different in this sample (ANOVA *P*=0.69).

**Figure 1:**
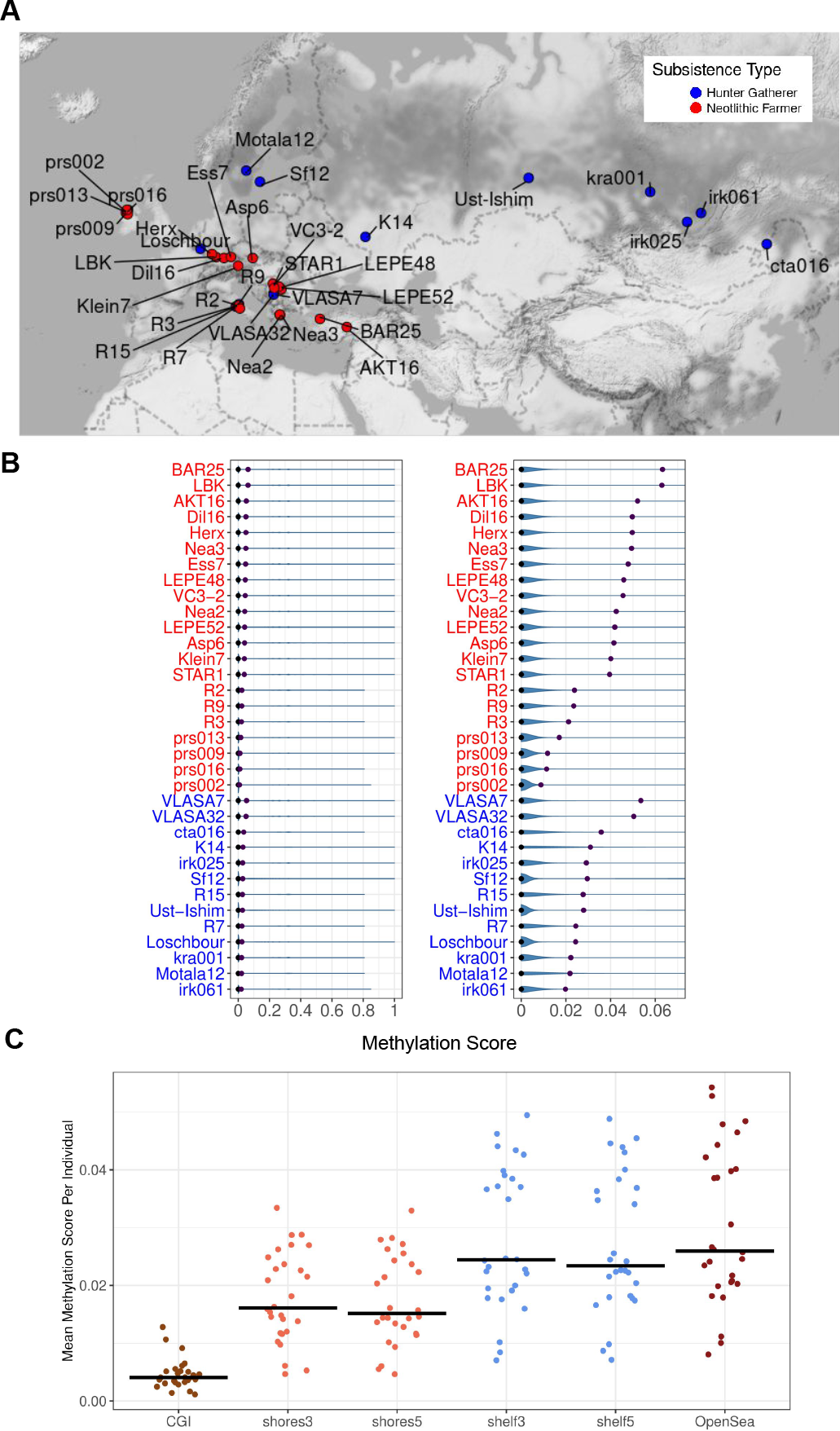
(A) The excavation locations of the ancient individuals included in this study. Colour coding indicates subsistence type. (B) Left panel: Violin plots of the methylation score (MS) data related to ancient individuals included in this study. The brown and blue points indicate the mean and the median, respectively. The x-axis shows the log2-transformed MSs. The y-axis represents the ancient individuals. Right panel: Zoomed-in version of the left panel. (C) The distribution of mean MS per individual on CpG islands and genomic sites representing shelves, shores and open seas. We used the R (Wickham, 2016) functions “ggmap” and “ggplot” for plotting geographical distributions and the CpG distributions.

To measure methylation rates, we used 13,270,411 autosomal CpG positions in the reference genome excluding variable positions (Methods). In this set an average of 9,238,400 (2,849,025-12,427,015) CpG’s were covered by at least one read per genome. Filtering for a minimum depth of 4 left us with an average of 3,006,714 (10,642-11,721,229) CpG positions per genome. Running epiPALEOMIX (Hanghøj *et al*., 2016) on this data, we computed the number of likely methylated (deaminated) and possibly non-methylated (not deaminated) reads, and the resulting methylation score (MS) for each CpG position per genome. The distribution of the MS values per CpG site across all 34 genomes revealed average methylation rates <7% (Figure 1B). This is much lower than the average CpG methylation rates in human tissues (60-80% (Anastasiadi *et al*., 2018; D. and Meissner, 2013)), but in line with published estimates from other paleogenomes (Hanghøj *et al*., 2016; Gokhman et al., 2014), and is caused by the indirect nature of methylation level measurements. We also observed multiple-fold differences in mean MS among the 34 paleogenomes (c.1% versus c.6%), which likely reflects technical effects rather than biological signals (Supplementary Tables 1-2).

Despite this variability, we found that CpG islands (CGIs), which are normally hypomethylated regions of the genome, show significantly lower MS scores (Wilcoxon signed rank test *P*<1e-10; Supplemental Table 3) across these 34 paleogenomes, compared to CGI shores (2 kb from CGI) and CGI shelves (4 kb from CGI) and more distant “open sea” areas (Figure 1C; Supplementary Figures 3-4). This indicates that the genome-wide MS values measured here have some degree of biological relevance.

### Tests for subsistence type, tissue and sex effects: few or no genes with evidence for systematic methylation differences

We next tested for differentially methylated genes (DMGs) related to subsistence type, tissue of origin (tooth or bone), and genetic sex. Before running the tests, to avoid possible biological effects being confounded by inter-genome variability in average MS values (Figure 1B), we normalized the dataset by subsampling reads for every individual genome randomly so that each genome gained a genome-wide mean MS of 0.02 (Methods). We performed this subsampling 20 times, creating 20 normalized replicate datasets. Using each of these replicates separately, and for each gene, we ran linear mixed effects models: all MS values across a gene as the response, subsistence type, tissue, and sex were fixed effects, and “individual” was the random effect.

We thus tested 9,657-9,660 genes across the 20 normalized datasets, with a median of 261 CpG positions in each gene (1-18,097). A total of 55-71 genes (0.5%-0.7% of tested genes) had ANOVA *P* < 0.05 after Benjamini-Hochberg (BH) correction for multiple testing for only subsistence type. The number of BH-corrected significant genes for tissue type and genetic sex were 19-39 (0.2%-0.4%) and 0-12 (0%-0.1%), respectively. Figure 2A shows the top genes identified for each factor. We note that this approach may be overestimating effects due to some degree of pseudoreplication, which we address below (Methods).

**Figure 2:**
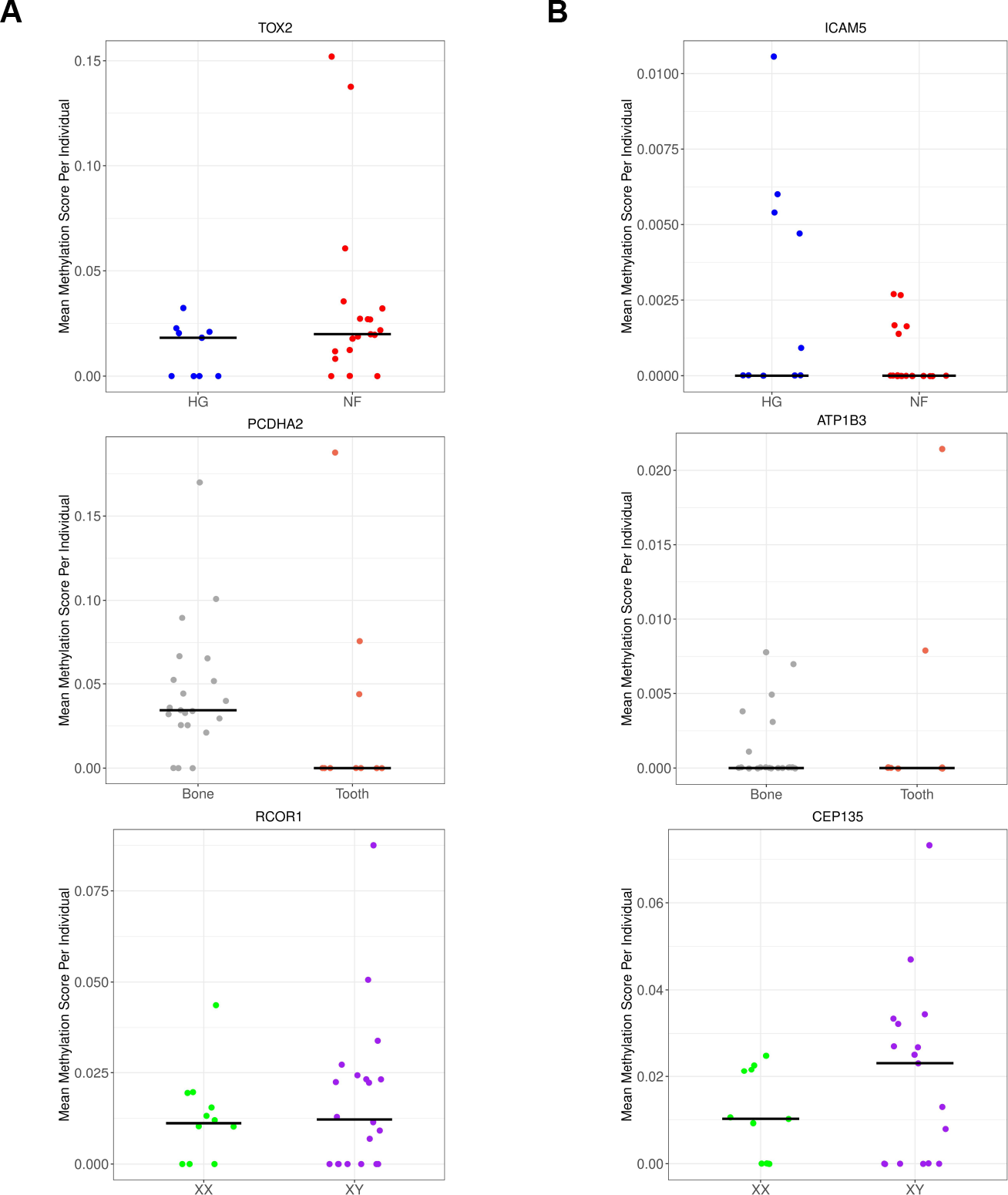
Representative genes with the most significant differential methylation results in linear mixed model analyses, related to subsistence type, tissue and sex, from top to bottom. The x-axis represents the factors while the y-axis represents the mean MS values per gene per individual. (A) Genes chosen using models with “individual” as random factor. Upper panel: *ICAM5* (subsistence type *P* < 0.01). Blue: HG; red: NF. Middle panel: *ATPB1* (tissue type *P* < 0.01). Coral: tooth; grey: bone. Lower panel: *CEP135* (genetic sex *P*=0.02). Green: female; purple: male. (B) Genes chosen using models with “laboratory-of-origin” as random factor. The color coding is the same as panel A. Upper panel: *TOX2* (subsistence type *P*=0.006). Middle panel: *PCDHA2* (tissue type *P*=0.03). Lower panel: *RCOR1* (genetic sex *P*=0.04).

We further performed enrichment analysis in Gene Ontology (GO) categories to identify possible functional roles of DMGs (those passing BH-corrected ANOVA *P*<0.05) relative to the background set of 9,657-9,660 genes across the 20 subsampled datasets. Although the most enriched GO terms included development- and regulation-related mechanisms (results for two randomly chosen datasets are shown in Supplementary Figures 4-5), none were significantly enriched after multiple testing corrections (BH-corrected Fisher’s exact test *P*<0.05).

We next repeated the previous analysis but this time using the “laboratory-of-origin” as random effect (instead of “individual”). The numbers of genes with sufficient information to execute ANOVA to compute P-values for all categories were 8,867-8,891 across the 20 subsampled datasets (Methods). This time, either no gene or a maximum of 2 genes were significant at BH-corrected *P* < 0.05 for any of the three fixed factors. The top genes are shown in Figure 2B; similar to those in Figure 2A no strong effects are visible even among these genes.

Instead of using the full data, summarizing MS values per gene might reduce noise and clarify the signal. For each of the 9,956 genes and all 34 individuals, we calculated the average MS across all CpG positions covered with a minimum of 4 reads per gene and averaged these across all 20 subsampled datasets (Methods). Using this dataset we performed a multi-dimensional scaling (MDS) analysis on Euclidean distances between individuals. This revealed that the K14 and Motala12 genomes, which also had the lowest coverage of CpG sites in our set, also behaved as outliers in their paleomethylome profiles (Supplemental Fig. 6). Removing these two genomes, an MDS plot of distances among the remaining 32 genomes revealed salient clustering by laboratory-of-origin (Figure 3).

**Figure 3:**
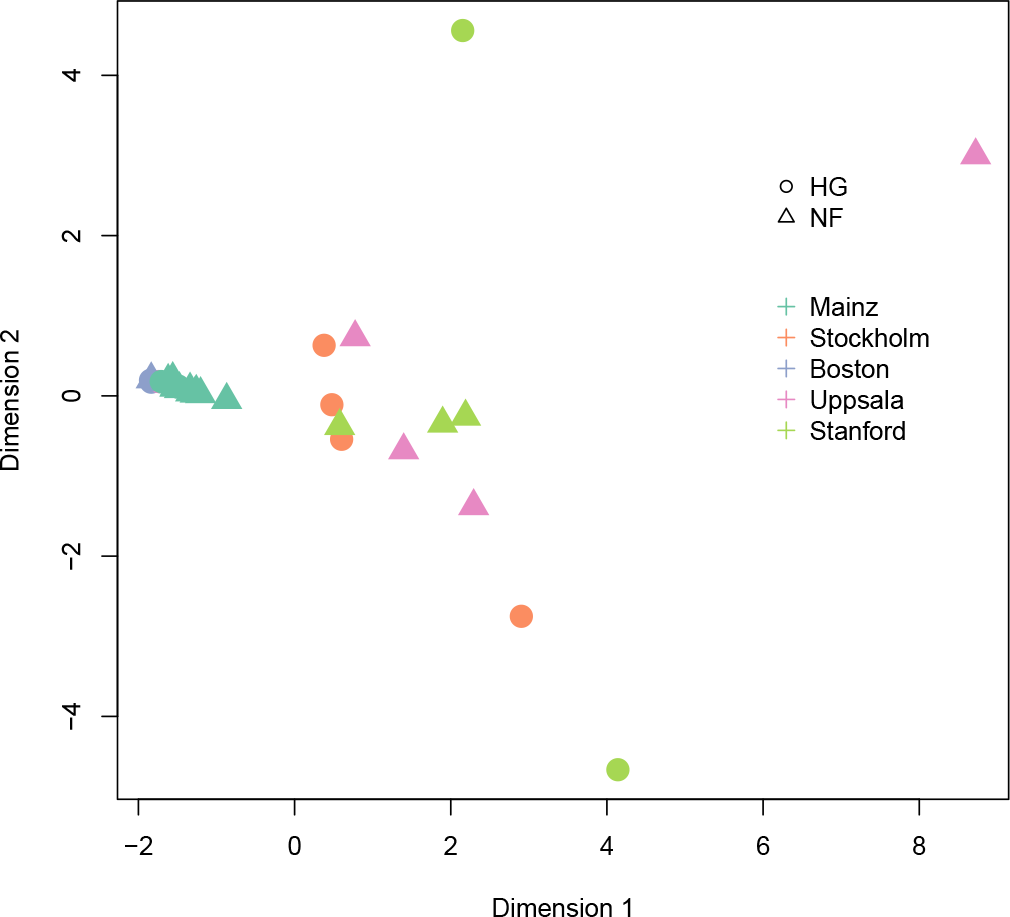
Multi-dimensional scaling (MDS) plot of 32 paleomethylome profiles. The data was created by Monte Carlo normalizing MS values followed by summarizing per gene. Circles: HG; triangles: NF. The colouring of the points represents the laboratory-of-origin of the samples (indicated by their city), as shown in the legend. The Motala12 and K14 genomes were not included in the analyses due to their outlier profiles compared to the rest likely representing technical effects (Supplementary Figure 6), which leaves us with five laboratories.

We further limited the dataset to 9,273 genes observed in a minimum of 20 individuals, and ran Kruskal-Wallis with laboratory-of-origin as an explanatory factor, excluding Motala12 and K14 individuals: we found an effect across 14% of genes tested (BH-corrected *P* <0.05). In contrast, running the same test using subsistence type, tissue, or sex as explanatory factors yielded no significant genes at this cutoff. Performing this analysis by limiting the dataset to a minimum of 25 or 30 individuals, using only genomes with 10x coverage, or using ANOVA produced qualitatively the same outcome. Hence, the laboratory-of-origin has a dominant signal in the data, which may obscure any biological effects.

### No significant correlation with subsistence-type effects in modern-day Africa

Although our previous analyses did not yield any clear signs of subsistence-related differential methylation, weak but authentic signals might still be detected by comparing our MS data with subsistence-related DMGs identified in modern-day populations, assuming Neolithic shifts would create convergent methylation signatures. We also decided to run this comparison both on our full dataset of HG-NF differences, but also separately on three paleomethylome datasets from different laboratories where both subsistence types were represented (Figure 4); we considered that this might help remove confounding between real signals and technical effects.

**Figure 4:**
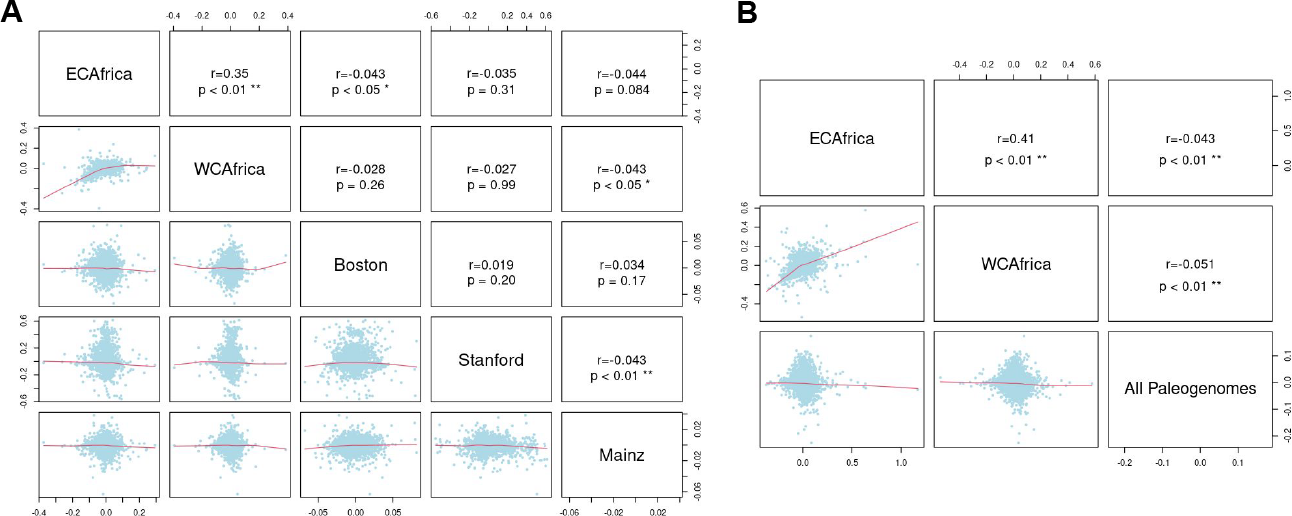
Pairwise comparisons of HG-agriculturalist DNA methylation differences between ancient and present-day human datasets. The lower triangle panels show the scatterplots of HG-agriculturalist average DNA methylation differences per gene in two datasets. The dataset and laboratory names are given on the diagonal panel. The upper triangle panel reports Spearman rank correlation coefficient *r* and *P*-values. “ECAfrica” and “WCAfrica” represent present-day HG-agriculturalist methylation differences (log fold-change) measured from humans in East Central and West Central Africa, respectively. (A) “Boston”, “Stanford” and “Mainz” stand for MS differences between HG-NF groups measured using only paleogenomes produced in the respective city (Supplementary Table 1). (B) “All Paleogenomes” stands for MS differences between HG-NF groups measured using all 32 paleogenomes (excluding Motala12 and K14).

To this end, we utilized methylation differences documented between modern-day HGs and agriculturalists in Central Africa, measured in whole blood samples using bisulphite treatment and the Illumina 450K array (Fagny *et al*., 2015). The authors of this study reported c.9000 and c.6000 genes that included CpG sites differentially methylated between independent groups of traditional HGs and agriculturalists living in Eastern Central Africa (EC Africa) or in Western Central Africa (WC Africa), respectively.

There were 7890 genes overlapping between our paleomethylome dataset and the modern African dataset. Across these genes, we calculated the correlation between methylation differences between HG-NF groups in our dataset, and the log-transformed mean fold change [log(FC)] values between modern-day HGs and agriculturalists groups in the EC Africa and WC Africa datasets as calculated by Fagny and colleagues (Methods). We observed a significant positive correlation (Spearman’s rank correlation coefficient *r*=0.34, *P* <0.01; Figure 4) between methylation differences in modern-day HGs and agriculturalists measured in EC Africa and WC Africa, in line with the original publication (Fagny *et al*., 2015). However, no consistent correlation could be observed between HG-NF differences within our paleomethylome dataset or between differences in the paleogenomes and HG-agriculturalist differences measured either in EC Africa or in WC Africa. Two nominally significant correlations were in fact negative and all correlations were weak (Spearman’s *r*<0.05) across 2507 genes (Figure 4).

### Lack of X chromosome methylation signatures among the 34 paleogenomes

The X chromosome (chrX) is expected to be methylated at higher rates in females compared to males, due to female X inactivation (Liu *et al*., 2010). Indeed, Liu and colleagues recently reported clear clustering of X chromosome MS values measured in ancient female and male horses (Liu *et al*., 2023). To investigate such signal among the 34 paleogenomes, we prepared a chrX paleomethylome dataset using the same steps as before, including normalizing by randomly subsampling once to 0.02x mean MS score. We tested each chrX gene for sex differences using ANOVA, using either “laboratory-of-origin” or “individual” as random factors. Unexpectedly, no chrX gene was significant for sex after the BH correction (*P* >0.05). A plot of chrX MS distributions between female and male individuals across all 34 paleogenomes, or only using 13 NF paleogenomes from Mainz, similarly revealed no obvious difference between sexes (Figure 5). This suggests that the overall biological signal in the dataset is indeed limited.

**Figure 5:**
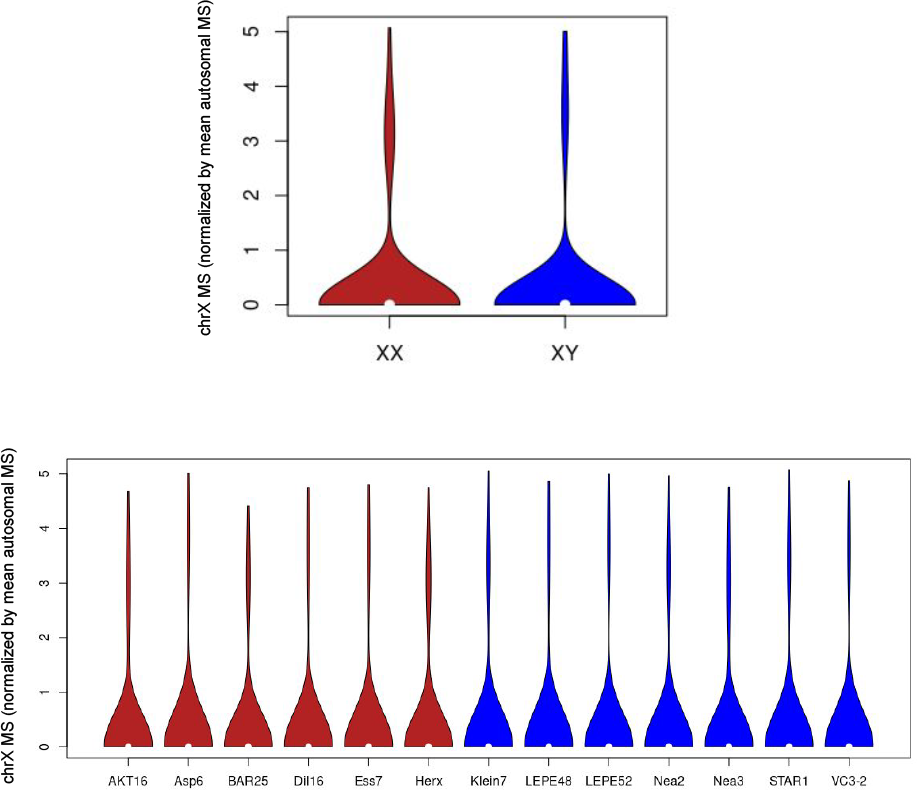
Violin plots of chrX MS values divided by the mean autosomal MS for that same individual Upper Panel: chrX MS values of all 34 paleogenomes. Lower Panel: chrX MS values from NF individuals from Marchi and colleagues (Marchi *et al*., 2022) These were chosen to remove the possible influence of other factors (different laboratory and subsistence type effects) on chrX methylation estimates.

## Conclusion

Today, thousands of human paleogenomes are being produced every year and there is growing interest in using these to study biological processes beyond historical and social questions (Orlando *et al*., 2015). This includes the study of DNA methylation levels. Even though cytosine methylation appears to survive in aDNA (Pedersen *et al*., 2014; Gokhman *et al*., 2014; Seguin-Orlando et al., 2015), it has been yet unclear whether the highly variable nature of the published paleogenomic data could allow reproducible signals to be inferred from joint datasets from different laboratories.

Here we investigated biological signals related to tissue source, sex, as well as subsistence type in a heterogeneous paleomethylome dataset comprising genomes from 6 different laboratories. We limited the calls to CpG sites with a minimum of 4 reads, normalized the data by subsampling to account for average coverage differences, and ran analyses using a number of different comparative approaches. Beyond hypomethylation of CGIs, we were unable to recover any biological signal that reached genome-wide statistical significance.

Whether universal subsistence-type effects related to hunter-gatherer versus agriculturalist lifeways might be prevalent in bone methylomes is an open hypothesis. Hence, not finding a consistent signal in this dataset may be not surprising and attributable to a diversity of possible effects, including the lack of a real convergent signal, or small sample sizes. However, the lack of tissue (bone versus tooth) or sex signatures, including on chrX, was unexpected.

Our negative results appear to contrast with the recent report by Liu and colleagues who identified systematic methylation signatures of sex, age, and castration in paleogenomes (Liu *et al*., 2023), or those by Hanghøj and colleagues (Hanghøj *et al*., 2016) who clustered genomes based on tissue type. However, the first study used >5x coverage genomes produced in the same laboratory, and the second study used data from two laboratories and only >14x genomes.

In our heterogeneous dataset, the most prominent clustering was by laboratory-of-origin. Such technical effects on methylation scores could be due to differences in mean depth-of-coverage, as well as variable coverage patterns across paleogenomes, which, in turn, could be driven by laboratory protocol differences in aDNA isolation, library preparation, or sequencing. We hypothesize that such technical variability overshadows any differential methylation signals that are subtle and measured indirectly. Hence, strict control of technical effects and the use of relatively high coverage (e.g. >5x) genomes appears to be a minimum requirement for future paleomethylome studies.

## Methods

### Genome data selection and preprocessing

We selected UDG/USER-treated shotgun-sequenced genomes from published genomic data including including 13 HGs and 21 NFs from West and North Eurasia. Sample-related information can be found in Supplementary Table 1. We note that the Siberian Bronze Age individuals were included in the HG category since these groups had an HG-like lifestyle with a diet composed mainly of marine and freshwater products (Kilinç *et al*., 2021). We chose to limit our sample to West and North Eurasia in order to limit the effect of differences in population genetic background but also tried to keep our sample large enough to increase power. We used the R (Wickham, 2016) function “ggmap” for plotting the chosen individuals’ geographical distributions (Figure 1A).

All data was downloaded as BAM or FASTQ files from the European Nucleotide Archive (ENA; https://www.ebi.ac.uk/ena), with reference numbers listed in Supplementary Table 1. All FASTQ and BAM files were remapped on *Homo sapiens* genome assembly hs37d5 using bwa aln with parameters “-l 16500 -n 0.01 -o 2” (Li and Durbin, 2009). We filtered out reads of size less than 35 bps, with a mapping quality (MAPQ) of less than 30, and with more than 10% mismatches to the reference genome. We verified the effectiveness of the UDG/USER treatment by studying the PMD profiles created using “pmdtools” (Skoglund *et al*., 2014) on each genome (Supplemental Figure 1, 2).

We called all CG dinucleotide autosomal positions (n= 26,752,702) from the human (hg19) reference genome using the R Bioconductor package “BSgenome.Hsapiens.UCSC.hg19” (Pagès, 2019) and stored these in a BED file. We then filtered these by removing any positions that overlapped with SNP positions from dbSNP 142 (Sherry *et al*., 2001). Our aim here was to avoid confounding between methylation signals and real variants at CpG positions. There remained 13,270,411 autosomal CpG positions in the reference genome.

We downloaded CpG island (CGI) positions for hg19 from the UCSC Genome Browser (Karolchik *et al*., 2004). We termed 2 kb sequences flanking CpG islands “shores” (upstream regions “shores5” and downstream regions “shores3”), 2 kb sequences flanking the shores “shelves” (upstream regions “shelves5” and downstream regions “shelves3”), and distal sites outside the CpG island regions as “open sea”, following (Hanghøj *et al*., 2016). “shores3” shores5

### Methylation score calculation

We chose to use the software epiPALEOMIX (Hanghøj *et al*., 2016) over DamMet (Hanghøj *et al*., 2019); the latter is an alternative methylome mapping software developed by the same group but is described as requiring ≥20x coverage to generate reliable results. Since our dataset median was much lower we decided to use epiPALEOMIX. epiPALEOMIX requires UDG/USER-treated and ≥2x-coverage genomes (we still included three genomes <2x to increase our sample size). The BAM file, the hg19 reference fasta file, the reference BED file for CpG positions and the library type of the sample (single-stranded/double-stranded) were given as input. We thus constructed our sample set and epiPALEOMIX input files according to these criteria.

We filtered the epiPALEOMIX output files for each CpG position having ≥4 reads to increase the precision of the MS values. We further generated a file that included the information related to the chromosome number, CpG position, and the MS values of each ancient individual as a column by joining all the files by CpG positions. Missing values were presented by “NA”.

We calculated average MS values per CpG position for each individual from the epiPALEOMIX outputs. Let *n*_1*i*_ denote the number of deaminated reads and *n*_0*i*_ denote the number of non-deaminated reads in genome *i*. We then calculated: 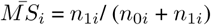. We also plotted the MS values per individual (Figure 1B) using ‘ggplot2’ function in RWickham (2016).

We performed gene annotation using the UCSC Genome Browser table for the hg19 assembly containing only exons (Karolchik *et al*., 2004). After that, we calculated MS at the promoter sites (4 kb long) by using 2kb upstream of the first exon on the positive strand.

We also ran epiPALEOMIX on the X chromosomes (chrX) of the same 34 individuals. These chrX datasets were prepared employing the same steps used in the autosomal datasets.

### Monte Carlo normalization

Given the large differences in mean MS values among the genomes (Figure 1), we normalized our ANOVA dataset, which includes all the reads corresponding to CpG positions per individual, by random subsampling the reads so that every individual in the dataset has 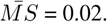. Note that here we again only use CpG positions with ≥4 reads in each genome. We chose 0.02 as a target as this was on the lower end of our 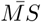 distribution.

Let *n*_1*i*_ denote the number of originally deaminated reads in genome *i*, and let *n*_0*i*_ denote the number of originally non-deaminated reads in the same genome. We proceeded as follows: a) If genome *i* had original mean MS < 0.02: we subsampled from *n*_0*i*_ a random subset *n*_0*is*_ as *n*_0*is*_=49*n*_1*i*_, so that *n*_1*i*_/ (*n*_0*is*_ + *n*_1*i*_) = 0.02. b) If genome *i* had original mean MS > 0.02: we subsampled from *n*_1*i*_ a random subset *n*_1*is*_ as *n*_1*is*_=*n*_0*i*_/49, so that *n*_1*is*_/ (*n*_0*i*_ + *n*_1*is*_) = 0.02.

We ran random resampling using the function “sample” offered by R.

We repeated the random subsampling 20 times independently to produce 20 normalized datasets. The chrX dataset was also normalized in the same manner, separately. We note that normalization is performed using all reads (on autosomes, or chrX), not just ones that overlap genes. We also normalized the chrX dataset over the autosomal MSs and plotted violin plots for all CpGs and also for the Neolithic individuals reported by Marchi et al. 2022 using the function “vioplot” provided by base R (Figure 5).

### Gene methylation datasets

We used these 20 normalized datasets to compile methylation levels per gene, in two ways:

i. Full data for linear mixed models: Here we used all normalized MS values for all CpGs overlapping a gene. Each individual may be represented by multiple CpG positions per gene (median 261). We had 20 parallel subsampled datasets of gene MS values. Note that the numbers of genes and CpG positions in each of these 20 datasets were slightly different because of random sampling of reads (e.g. genes with one CpG position might not be represented in some datasets).
ii. Gene-averaged data: This single dataset was produced by calculating, per gene, the means of all CpG MS values and averaging these across the 20 subsampled datasets. This yielded a simplified dataset with 9,956 genes x 34 genomes.

### Statistical tests

We used tests from the R “stats” package. All the tests were carried out two-sided unless otherwise indicated. We adjusted *P*-values for multiple testing using the Benjamini-Hochberg procedure using the R “p.adjust” function.

### Linear mixed effects models

. We applied these to the full data (i) described above, where multiple CpG positions per gene represent an individual. Since we had fixed (subsistence type, tissue, and genetic sex) and random factors (individual or laboratory-of-origin) in the settings, we decided to conduct linear mixed-effects models employing the R “stats” package “aov” function (R Core Team, 2020). We tested two models that differed in their random factors for each gene:

Model 1: deamination ∼ subsistence type + tissue type + genetic sex + Error(individual)

Model 2: deamination ∼ subsistence type + tissue type + genetic sex + Error(laboratory-of-origin)

Here the response variable “deamination” is a binary [0,1] variable that describes how many reads falling into each gene are deaminated or not. Note that this approach suffers from pseudoreplication, because the observations (reads) per locus are dependent when multiple reads map to the same locus. To overcome this, we also used the gene-averaged data (ii) described above. This time we applied ANOVA and Kruskal-Wallis tests on MS values per gene but without an individual component, using the R “stats” package “aov” and “kruskal.test” functions, respectively (R Core Team, 2020). Here we have a single observation per gene, and thus the results do not suffer from pseudoreplication.

### Multidimensional Scaling Analysis

We carried out multidimensional scaling (MDS) analysis on our gene-averaged dataset which included mean MSs per gene averaged 20 subsampled datasets. We used the R’s “cmdscale” function. We ran MDS both including all 34 individuals, or using 32 individuals after excluding extreme outliers Motala12 and K14 (Figure 3, Supplementary Figure 6).

### Gene Ontology Enrichment

Gene Ontology (GO) (Consortium, 2008) enrichment analysis (Subramanian *et al*., 2005) was performed by comparing gene sets with evidence for significant effects (for subsistence type, tissue type, or genetic sex) that had BH-adjusted P-values <0.05 from the linear mixed-effects models. We used the R “topGO” (Alexa and Rahnenfuhrer, 2019) and “org.Hs.eg.db” packages (Carlson, 2019) to collect GO information for the genes. The background gene sets included all 9,657-9,660 genes across the 20 normalized datasets included in the analyses. We ran the Fisher’s exact test within “topGO”, and used its “elim” algorithm for transversing the GO hierarchy (removing genes from significantly enriched lower nodes) (Alexa and Rahnenfuhrer, 2019). We also filtered the output to have ≥5 genes per GO term by using the “nodeSize” option while creating the GO data. The *P*-value threshold for the significance of the GO terms was chosen to be 0.01. We also visualized resulting GO terms using reviGO with default parameters (Supek *et al*., 2011). Results for two randomly chosen datasets (out of 20 datasets) are shown in Supplementary Figures 4-5.

### Subsistence Type-Related Methylation Differences in Ancient Eurasian vs Modern African Datasets

A recently published study uses blood samples taken from individuals to compare modern-day HG (MHG) and modern-day farmer (MF) blood methylation profiles in West and East African rainforests (Fagny *et al*., 2015). We used the results file of the study which contained the multiple-testing corrected *P*-values and the logarithm of methylation fold-change between MFs versus MHGs (logFC). In total, the dataset contained 365,401 CpG positions overlapping 19,672 genes. We used this information to estimate correlations between our results and the modern results reported by the original study.

We tested the co-directionality between the logFC values in this dataset and NF-HG differences we calculated in our methylome dataset. In other words, we compared farmer versus HG differences in MS scores *δ*_*MS*_*F −HG* across overlapping genes between pairs of datasets. Given the variability of MS profiles among genomes from different laboratories, we performed this comparison using sub-datasets from 3 different laboratories that contained both NF and HG individuals (Boston, Stanford, Mainz; see Supplementary Table 1), and also using 12 HG and 20 NF genomes excluding Motala12 and K14 individuals. We calculated the Spearman’s rank correlation between *δ*_*MS*_*F −HG* values from two datasets across common genes using the R “stats” package function “cor.test” (R Core Team, 2020). We plotted the lowess regression lines for the main laboratory of origins using the R “graphics” package “pairs” function with the “panel.cor” and “panel.smooth” parameters (Figure 4). The correlations and *P*-values are calculated using Spearman’s rank correlation method. For plotting we used the R “graphics” package and “ggplot2” package functions (Wickham, 2016).

## Software Availability

## Acknowledgements

We thank past and present members of METU CompEvo, Arev Pelin Sümer, İdil Yet, H. Melike Dönertaş, Güls, ah Merve Kilinç, and Kivilcim Başak Vural for kind assistance and discussions. We also thank Maud Fagny and Lluis Quintana-Murci for kindly sharing data. This work was supported by the European Research Council (ERC) Consolidator grant “NEOGENE” (Project No.: 772390 to M.S.) (https://erc.europa.eu).

## Author Contributions

S.S.Ç., R.N.F., D.K., M.S.: conceived the study and analyses, analysed data, and wrote the manuscript.

