## Supplementary Information for "Investigating DNA methylation changes associated with food production using paleogenomes"

**Supplementary table legends**

**Supplementary Table 1.** Individual: individual genome ID as in the original article. Laboratory: the city where work has been conducted (or the senior author is based). Coverage: mean genome depth-of-coverage. Library: single or double-stranded library.	Country: the country of origin of the ancient individual. Subsistence type: hunter-gatherer (HG) or Neolithic farmer (NF). Tissue: the tissue sampled for aDNA; bone or tooth. Total CpG positions: CpG positions out of 13 million (after excluding variable sites) that are covered by minimum 1 read. Total filtered >= 4: CpG positions out of 13 million (after excluding variable sites) that are covered by minimum 4 reads. Mean MS: mean methylation score (MS) per genome.

**Supplementary Table 2.** Number of genomes and mean methylation scores (MS) across those genomes per laboratory.

**Supplementary Table 3.** Pairwise Wilcoxon rank sum test results (*P*-values) over CGI, “shores3”, “shores5”, “shelf3”, “shelf5”, and “open sea” regions (Methods). Here we calculated the mean MS values for the respective regions, and performed comparisons using the values of all paleogenomes (n=34).

**Supplementary figures**

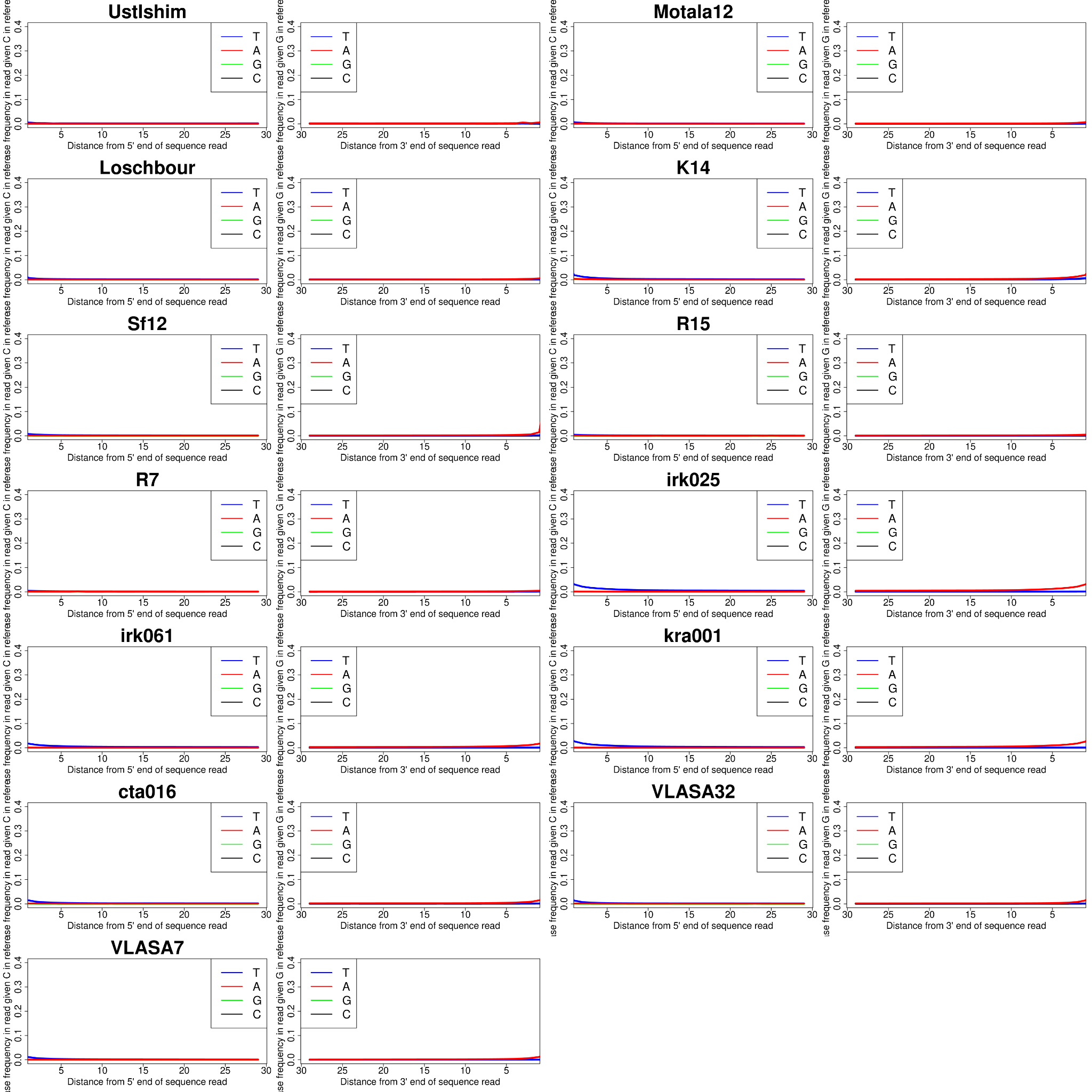

**Supplementary Figure 1.** Postmortem damage profiles of HG paleogenomes. The lack of excess C->T or G->A transition signals at 5’ and 3’ ends of reads, respectively, confirms the libraries were UDG-treated.

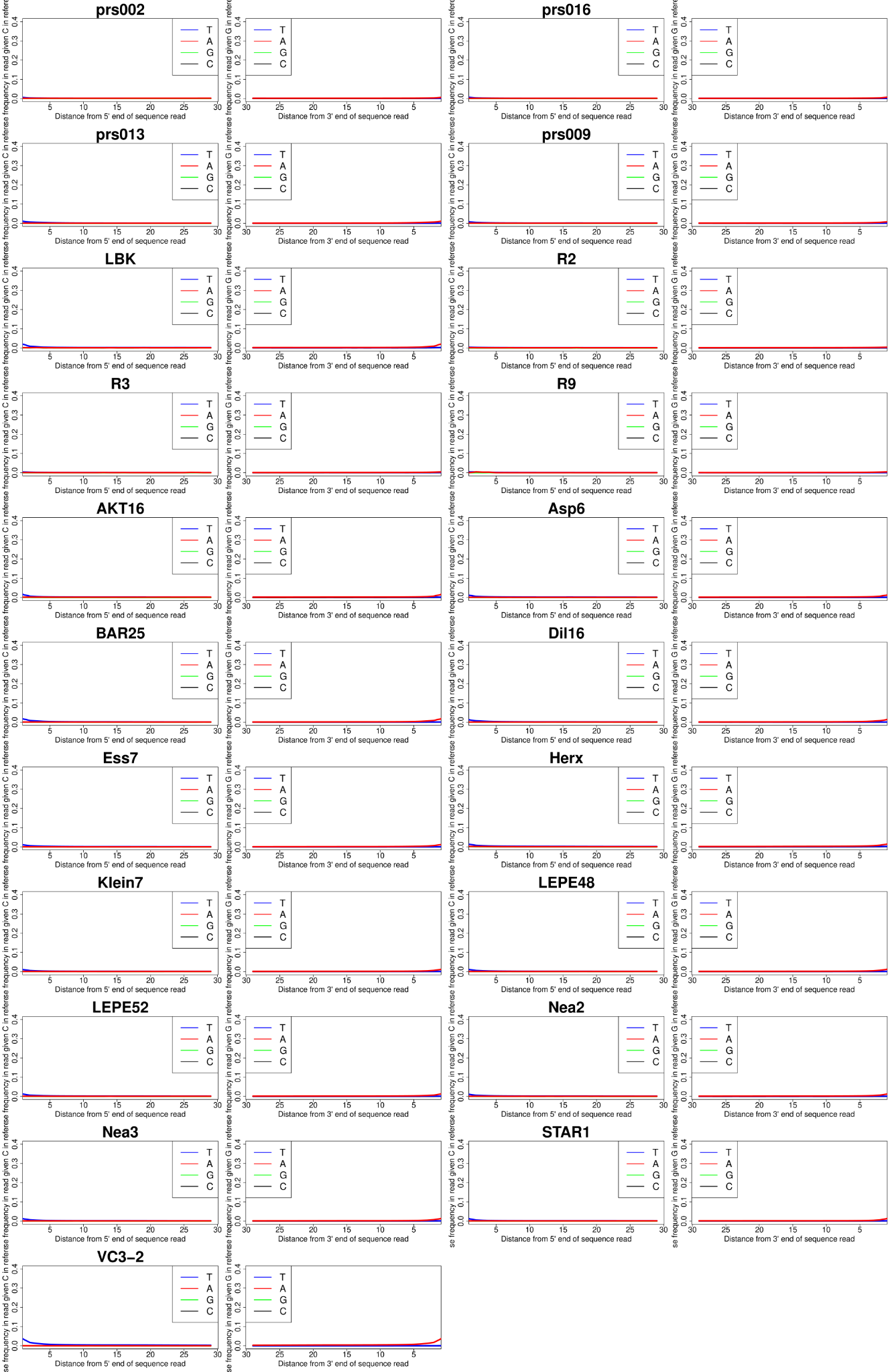

**Supplementary Figure 2.** Postmortem damage profiles of NF paleogenomes. The lack of excess C->T or G->A transition signals at 5’ and 3’ ends of reads, respectively, confirms the libraries were UDG-treated.

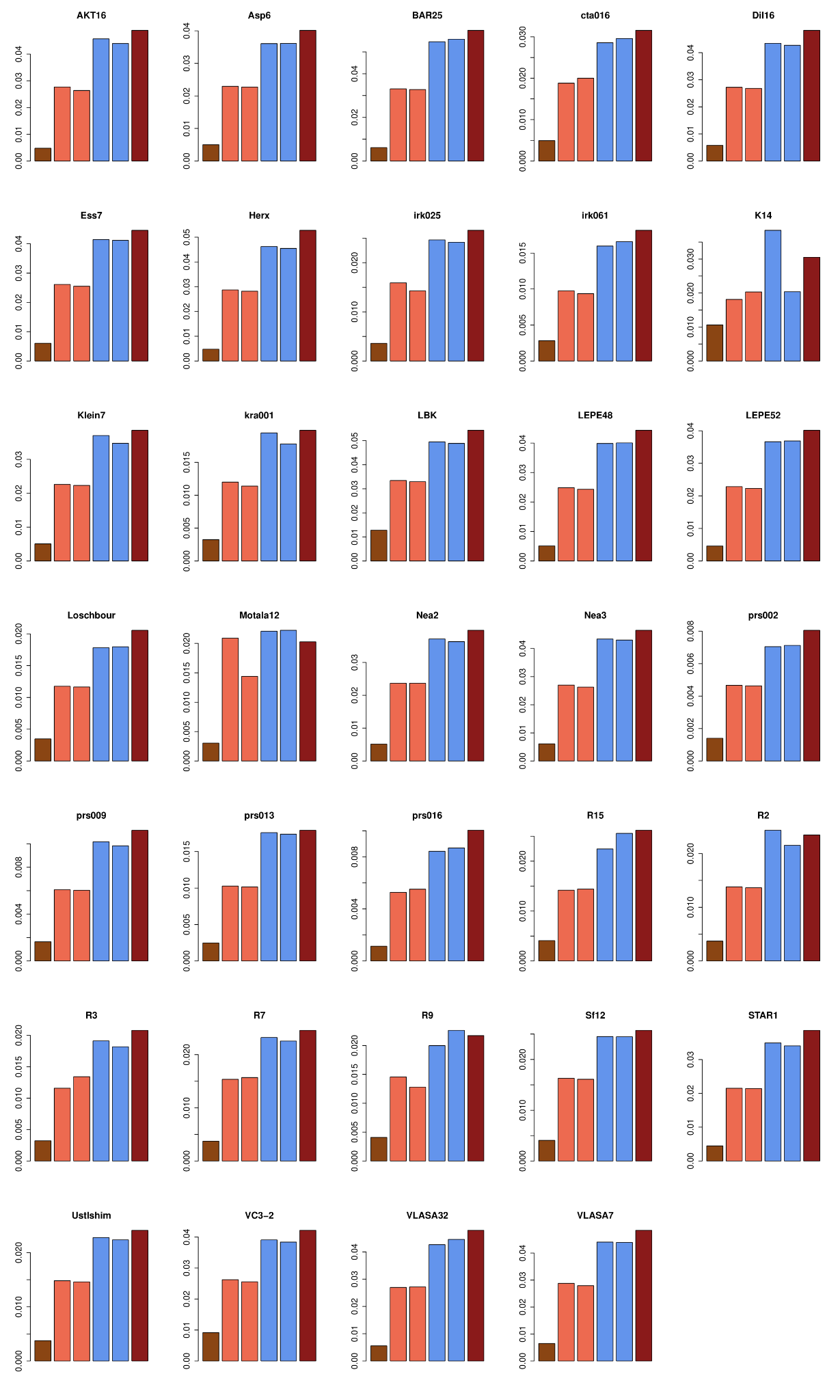

**Supplementary Figure 3.** The mean Methylation Scores (MS) on CpG islands (CGIs), “shelves”, “shores” and “open sea” areas of the genome per individual (continued in Supplementary Figure 4). The y-axes represent the mean MS and the x-axes indicate the genomic areas named above. The color brown represents CGIs, coral represents “shores”, blue indicates “shelves” and red represents “open sea” areas.

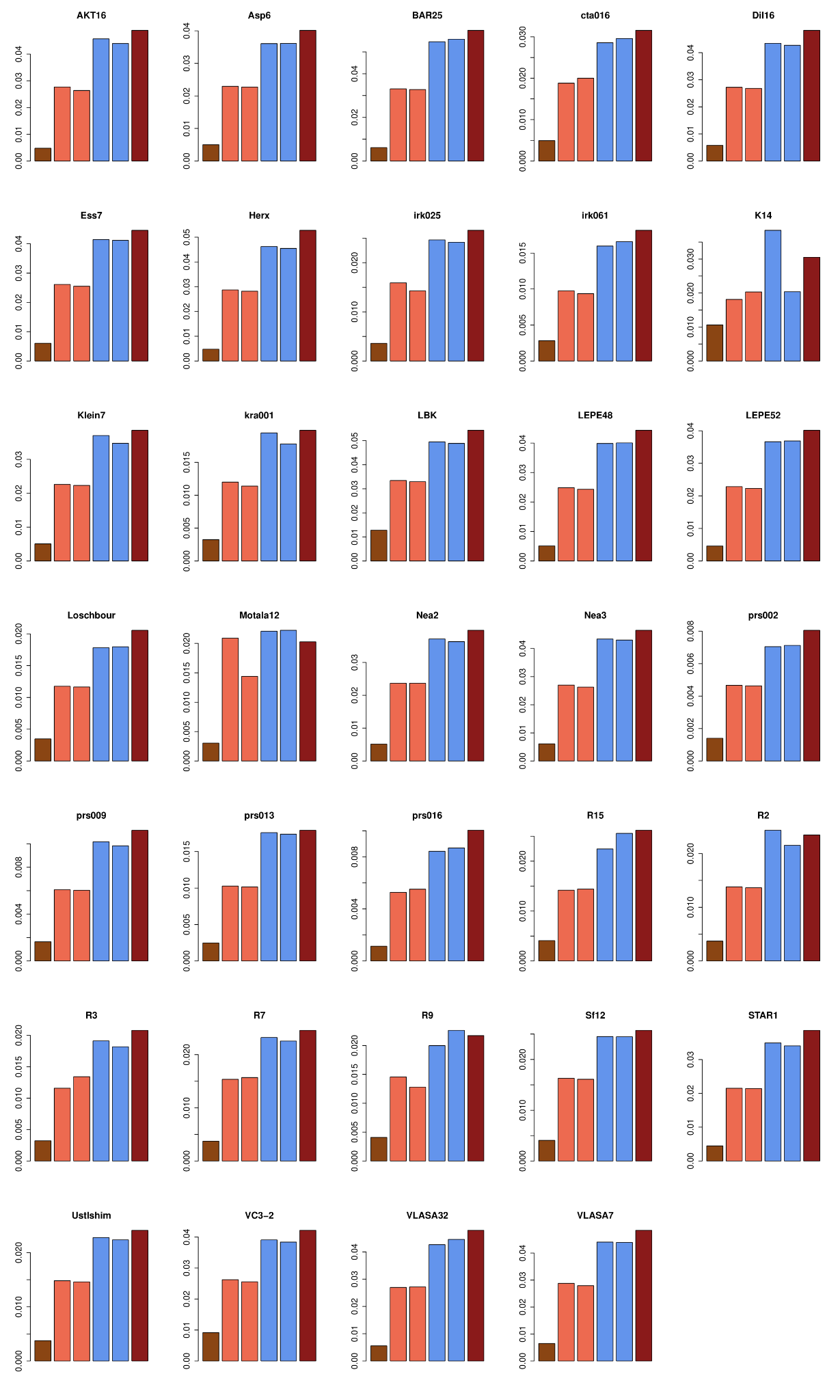

**Supplementary Figure 4.** The mean Methylation Scores (MS) on CpG islands (CGIs), “shelves”, “shores” and “open sea” areas of the genome per individual (continued from Supplementary Figure 3). The y-axes represent the mean MS and the x-axes indicate the genomic areas named above. The color brown represents CGIs, coral represents “shores”, blue indicates “shelves” and red represents “open sea” areas.

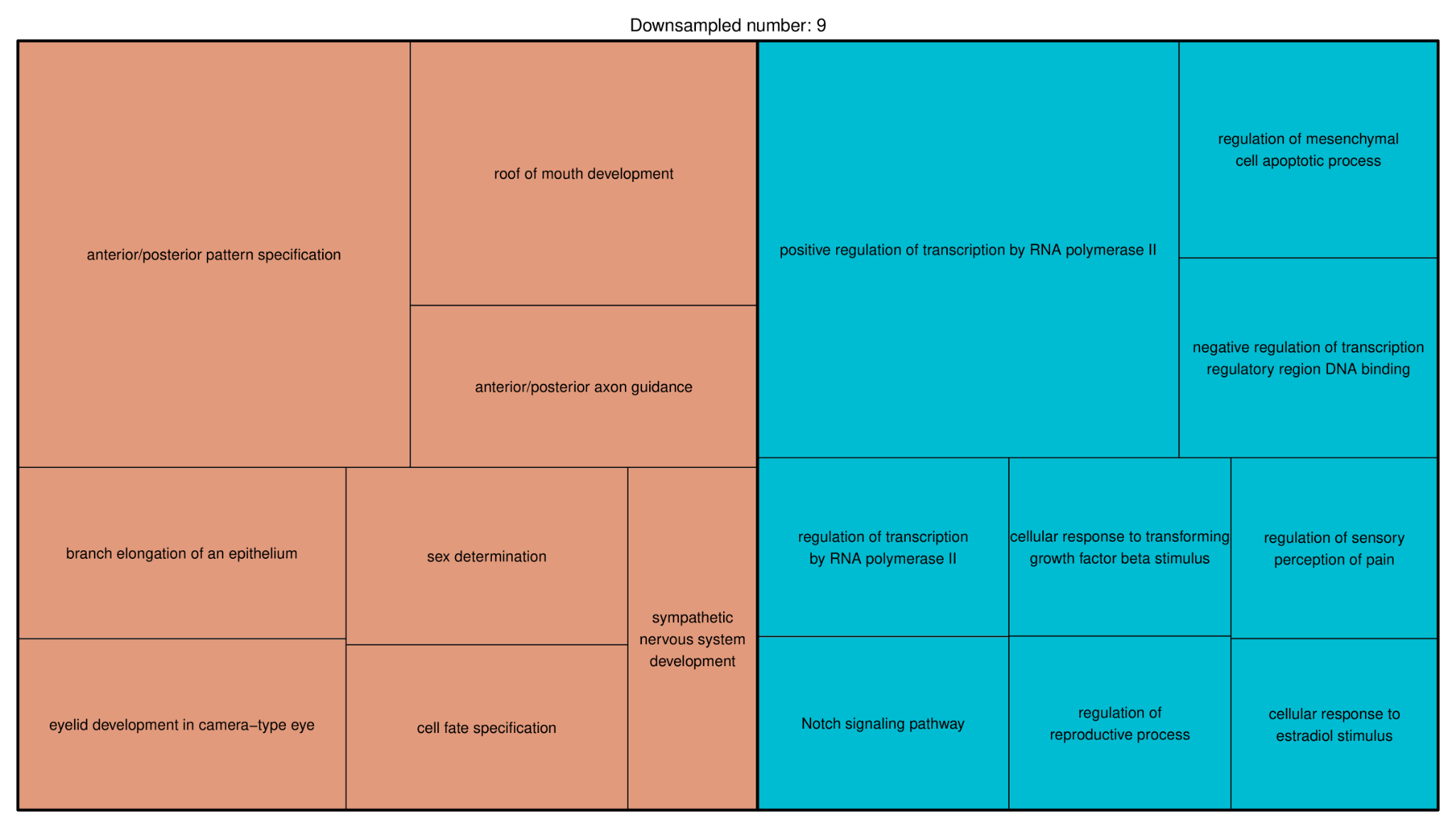

**Supplementary Figure 4.** Gene Ontology enrichment analysis results on subsampled dataset 9. Developmental processes are grouped on the left side (brown). Regulation processes are grouped on the right side (blue).

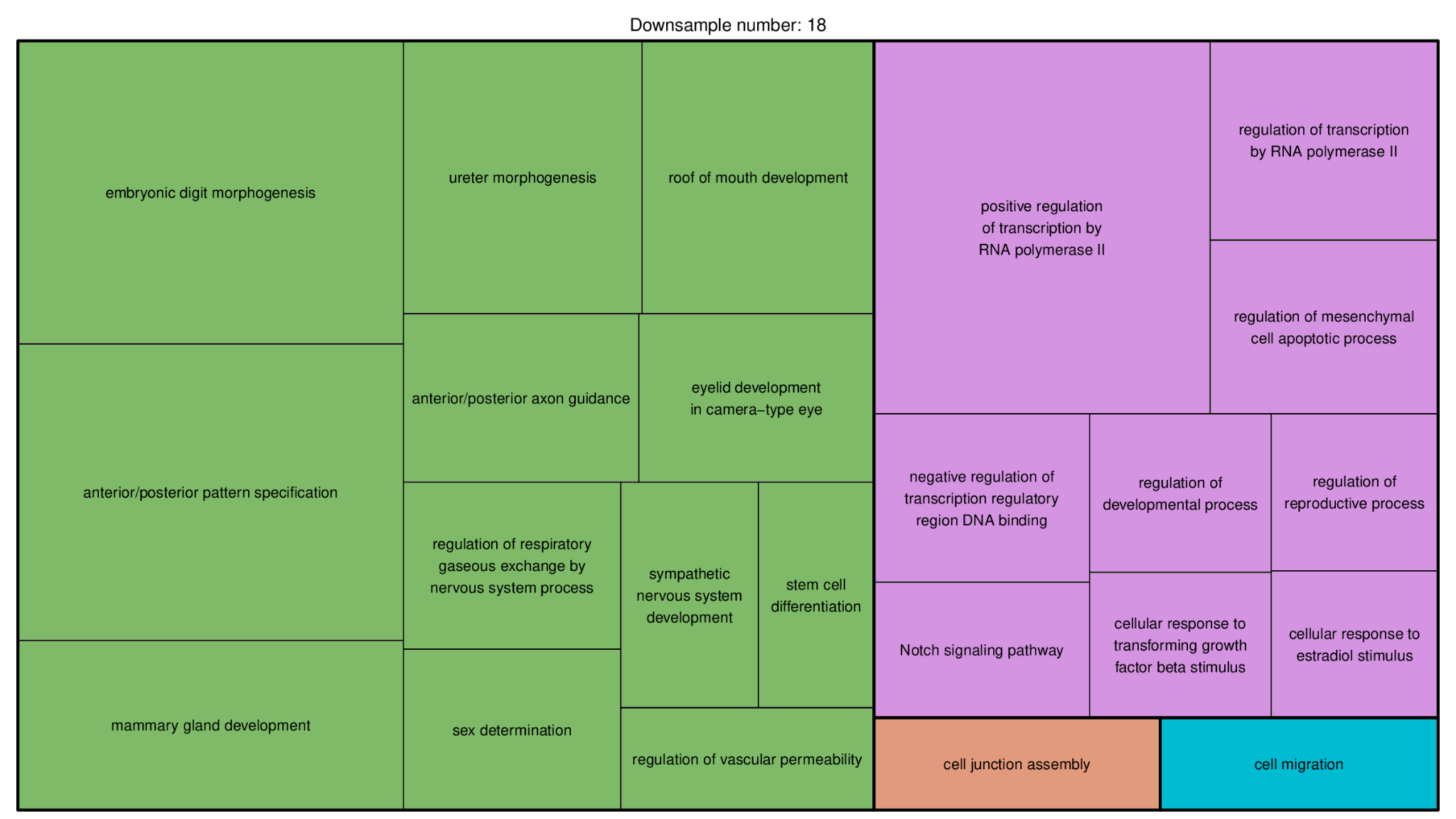

**Supplementary Figure 5.** Gene Ontology enrichment analysis results on subsampled dataset 18. Developmental processes are grouped on the left side (green). Regulation processes are grouped on the upper right side (purple).

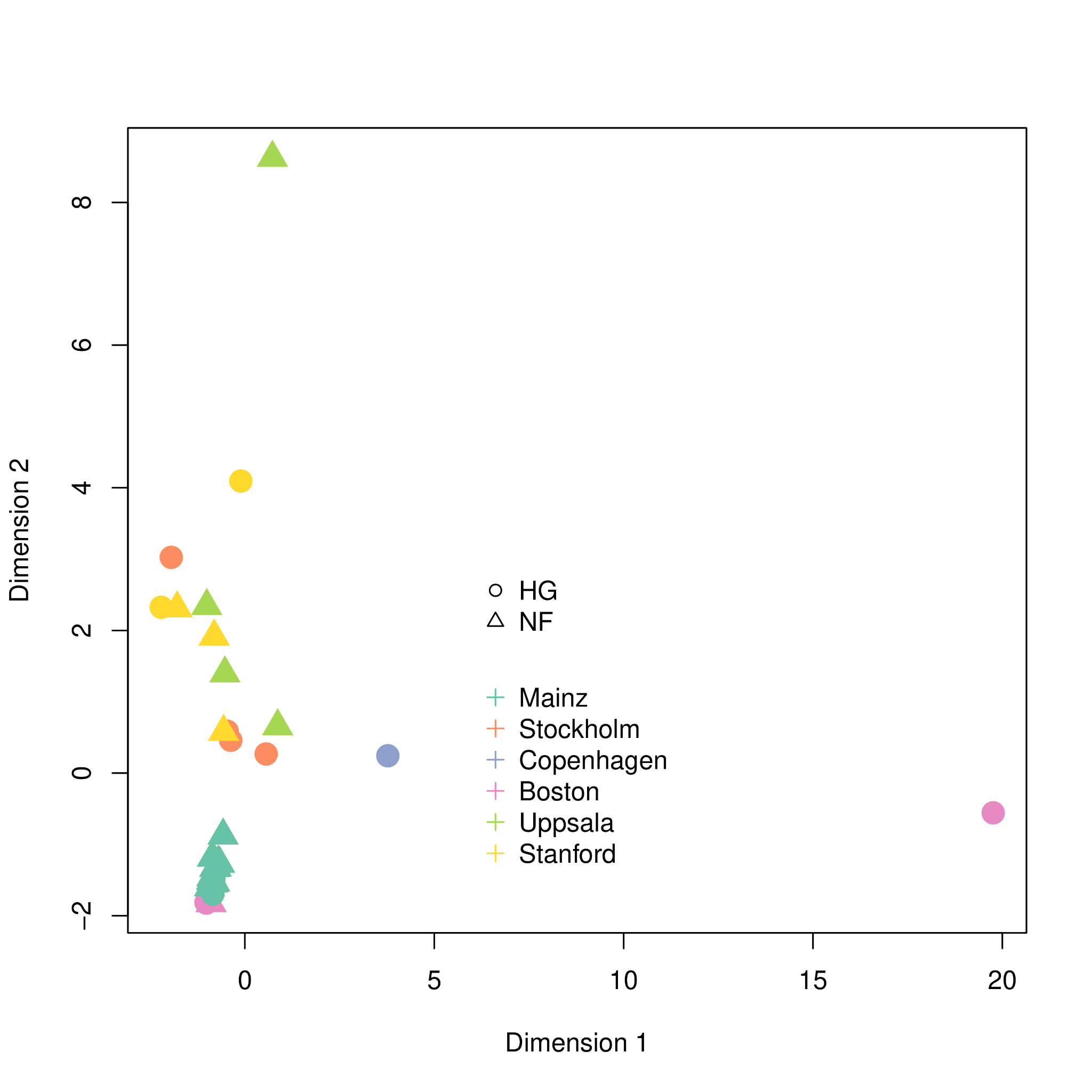

**Supplementary Figure 6.** Multidimensional scaling of the individuals by gene-averaged MS values. Hunter-Gatherers are indicated by the circles and Neolithic Farmers are represented by the triangles. Coloring of the illustration corresponds to the laboratory-of-origin of the samples. There are 6 different laboratories of which selected samples are produced from; Mainz (dark green), Stockholm (orange), Copenhagen (blue), Boston (pink), Uppsala (green), and Stanford (yellow). The Motala12 genome is on the right bottom and stands as an extreme outlier. The K14 genome the blue circle also shifted to the right of the main cluster.
